# The highly and perpetually upregulated thyroglobulin gene is a hallmark of functional thyrocytes

**DOI:** 10.1101/2023.06.23.546241

**Authors:** Simon Ullrich, Susanne Leidescher, Yana Feodorova, Katharina Thanisch, Jean-Baptiste Fini, Bernd Kaspers, Frank Weber, Boyka Markova, Dagmar Führer, Mirian Romitti, Stefan Krebs, Helmut Blum, Heinrich Leonhardt, Sabine Costagliola, Heike Heuer, Irina Solovei

## Abstract

Abnormalities are indispensable for studying normal biological processes and mechanisms. In the present work, we draw attention to the remarkable phenomenon of a perpetually and robustly upregulated gene, the thyroglobulin gene (*Tg*). The gene is expressed in the thyroid gland and, as it has been recently demonstrated, forms so-called transcription loops, easily observable by light microscopy. Using this feature, we show that *Tg* is expressed at a high level from the moment a thyroid cell acquires its identity and both alleles remain highly active over the entire life of the cell, i.e. for months or years depending on the species. We demonstrate that this high upregulation is characteristic of thyroglobulin genes in all major vertebrate groups. We provide evidence that *Tg* is not influenced by the thyroid hormone status, does not oscillate round the clock and is expressed during both the exocrine and endocrine phases of thyrocyte activity. We conclude that the thyroglobulin gene represents a valuable model to study the maintenance of a high transcriptional upregulation.

**SUMMARY STATEMENT:** The thyroglobulin gene is highly and permanently expressed in thyrocytes of all vertebrates, at any condition and round the clock, offering a unique model to study mechanisms of high upregulation maintenance and chromatin dynamics

## INTRODUCTION

It has been recently shown that highly expressed genes expand from their harboring loci and form so called transcription loops (TLs) (Mirny and Solovei, 2021). The expansion of highly expressed genes is attributed to their intrinsic stiffness acquired through decoration of the gene axis with multiple jam-packed RNA polymerases II (RNAPIIs) with attached nascent RNA transcripts (Fig. 1A). Therefore, not only the gene length but also the high gene expression level are essential for formation of a microscopically resolvable transcription loop (Leidescher et al., 2022). Exactly this combination of gene properties is extremely rare, especially in mammalian cells. Consequently, TLs had not been described until the recent serendipitous discovery of TLs formed by the unusually upregulated thyroglobulin gene.

**Figure 1.**
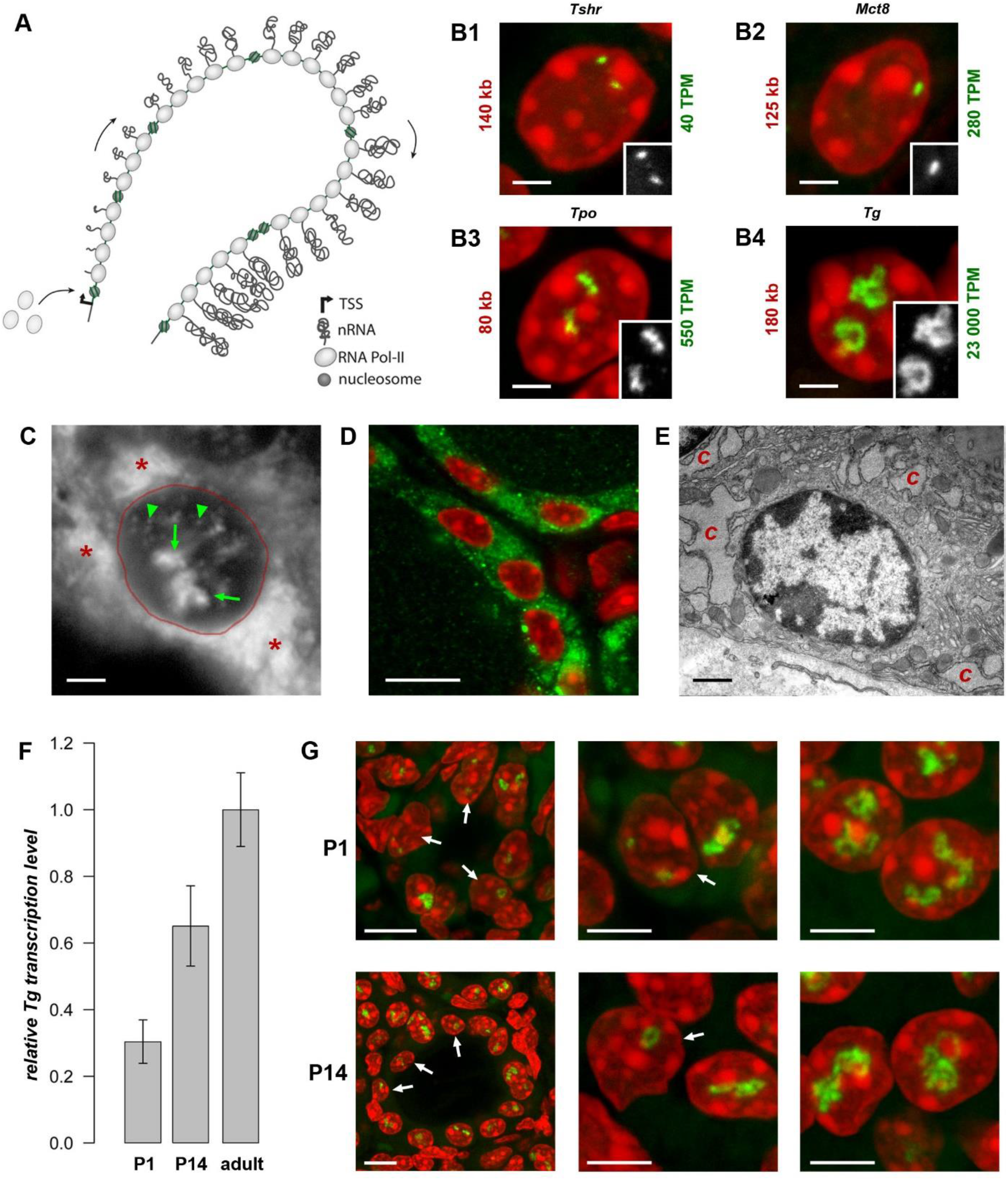
Transcription loops. **(A)** Schematics of a transcription loop (TL) formed by RNAPIIs moving along a gene and carrying nascent RNA transcripts. **(B)** Visualization of four thyroid-specific genes by RNA-FISH. In contrast to the lowly expressed genes *Tshr* and *Mct8* (B1, B2), the other two genes, *Tpo* and *Tg* (B3, B4), exhibit larger TLs with sizes correlated to their expression levels. Length and expression level of the genes are indicated on the left and right of the panels, respectively. For better comparison, TLs are shown as grey scale images in the insertions. Note, that in case of the *Mct8* gene located on X chromosome, there is only one signal, because the tissue originated from a male mouse. **(C)** FISH with an oligoprobe for *Tg* mRNA labels TLs (*green arrows*), single mRNAs in the nucleoplasm (*green arrowheads*) and in the cytoplasm (*red asterisk*). **(D**,**E)** Immunostaining of TG (*green*) highlights the cytoplasm of thyrocytes (D), which is densely packed with remarkably hypertrophic cisterns (*c*) of endoplasmic reticulum (E). **(F)** Histogram showing comparative *Tg* transcript levels in thyroids of young mice (P1 and P14) in comparison to adult mice. The qPCR values were normalized to the transcription level of the thyroid specific transcription factor Pax8. Error bars are SDM. **(G)** Examples of thyrocytes from P1 (*top*) and P14 (*bottom*) thyroids. The *left column* shows follicles formed by thyrocytes with various degree of *Tg* TL development; the *right and middle columns* show thyrocytes at a higher magnification with fully or underdeveloped *Tg* TLs. Arrows point at nuclei with underdeveloped loops. Note that in some nuclei only one allele is active. Images on B-D and G are projections of *3-5 µm* confocal stacks; FISH and immunostaining signals are *green*; nuclei are counterstained with DAPI (*red*). Scale bars: B, C, E, *2 µm*; D, *70 µm*; G, follicle overviews on the left,*10 µm*, zoomed in nuclei, *5 µm*.

The thyroglobulin gene (*Tg*) is expressed exclusively in thyrocytes, which in vertebrates are arranged in follicles that, in turn, unite into thyroid glands in most vertebrate groups. The thyroglobulin protein (TG) accumulates in the follicle cavity, representing the major component of the so-called colloid, and serves as a long-storage precursor for synthesis of the thyroid hormones (THs), 3,3’,5,5’-Tetraiodo-L-thyronine (T4) and 3,3’,5’-Triiodo-L-thyronine (T3) (van de Graaf et al., 2001). Importantly, *Tg* is highly upregulated with a transcription level of ca. 23,000 TPM (Transcripts Per Million), exceeding by many fold the expression of other thyrocyte-specific genes, as well as housekeeping genes, e.g., ubiquitinase (6-fold), actin (9-fold), myosin (22-fold) and multiple ribosomal genes (from 10- to 20-fold). Presumably, *Tg* is transcribed in long transcription bursts separated by short infrequent pauses (Fig. 1A) (Leidescher et al., 2022).

The presence of two highly extended *Tg* TLs, corresponding to the two active gene alleles in more than 93% of nuclei (Leidescher et al., 2022), indicates that the gene is remarkably upregulated in almost every thyrocyte for a long time. Therefore, we were interested in investigating thyroglobulin gene activity more closely. In particular, we followed *Tg* activation in development and showed that the *Tg* TL is a hallmark of functional thyrocytes. We asked whether the thyroglobulin gene in other vertebrates is similarly upregulated and demonstrated that thyrocytes of all major vertebrate groups exhibit thyroglobulin TLs. We further investigated whether *Tg* is regulated by organismal or cellular physiological conditions and proved that *Tg* transcription and translation are not regulated by the thyroid hormone (TH) status and do not undergo circadian rhythmicity or intron retention, ruling out these possible mechanisms of temporal segregation of the exocrine and endocrine phases.

## RESULTS & DISCUSSION

### *Tg* is an exceptionally highly expressed gene in thyrocytes

In agreement with its length (180 kb in mouse) and level of transcription (23,000 TPM), the *Tg* gene forms very prominent transcription loops (Fig. 1B4). To examine other long thyroid-specific genes, we selected thyroid peroxidase (*Tpo*) with a length of 78 kb, the TH transporting monocarboxylate transporter MCT8 (*Slc16a2*) that is 125 kb long and the thyroid stimulating hormone receptor (*Tshr*) with a length of 139 kb. The RNA signals of *Slc16a2* and *Tshr*, which have comparable length and ca. 100-500 times lower expression levels than *Tg*, are small and only slightly elongated. On the contrary, the much shorter *Tpo* gene, which is about 80 kb long but expressed at ca. 550 TPM, noticeably expands and forms short but distinctive transcription loops (Fig. 1B). This finding confirms that formation of microscopically resolvable transcription loops is dependent not only on the gene length, but also on the intensity of transcription (Leidescher et al., 2022).

The high transcription of the *Tg* gene is consistent with high production of the thyroglobulin protein (TG). The *Tg* transcripts comprise 2.5% of the entire mRNA pool in mouse thyroid, similarly to 2.6% shown for human thyrocytes (Pauws et al., 2000). In accordance, the probe for the *Tg* mRNA highlights not only *Tg* TLs and intranuclear mRNA, but also the whole thyrocyte cytoplasm (Fig. 1C). In agreement with this, the cytoplasm is brightly stained with an antibody against TG (Fig. 1D), and exhibits dilated cisternae of Golgi apparatus, as well as a remarkable hypertrophy of the rough endoplasmic reticulum, which occupies most of the thyrocyte cytoplasm (Fig. 1E). The above observations corroborate the statement that the *Tg* gene is abnormally upregulated and thyrocytes produce immense amounts of *Tg* mRNAs and TG protein.

### Thyrocytes exhibit *Tg* TLs from the onset of their differentiation

In mouse, the onset of folliculogenesis starts at E15.5 and the thyroid gland forms before birth (De Felice and Di Lauro, 2011). We aimed at estimating the *Tg* gene expression in early postnatal development and performed qPCR on thyroid glands dissected at stages P1 and P14. We showed that *Tg* expression increases during development, reaching a 3.3-fold rise in the adult mice compared to P1 pups (Fig. 1F). RNA-FISH revealed that although *Tg* TLs in most P1 and P14 thyrocytes are fully developed, ca. 20% and 12% of the cells, respectively, possessed small, undeveloped loops and often only one active allele (Fig. 1G, arrows). Apparently, these cells represent freshly differentiated or still differentiating thyrocytes. Although mitotic cells in the follicle epithelium are relatively infrequent, in both developmental stages, we observed several thyrocytes entering or exiting the cell cycle. In particular, we noticed that *Tg* TLs are manifested when most of nuclear chromatin is condensed: they appear in early G1, remain visible in thyrocyte nuclei until early-to-mid prophase and are withdrawn only in the very late prophase (Fig. S1). Thus, we conclude, that the *Tg* gene is expressed at a high level from the moment cells acquire thyrocyte identity.

### The *Tg* gene is highly expressed and forms transcription loops in all vertebrates

Initially, the thyroglobulin TLs have been described in mouse thyrocytes (Leidescher et al., 2022). However, THs are essential regulatory elements in the development and metabolism of all animals, and the mechanisms of TH generation are evolutionary conserved (Di Jeso and Arvan, 2016; Holzer et al., 2016). Therefore, we anticipated that in other mammals and other vertebrate classes, expression of thyroglobulin gene orthologues is also highly upregulated.

First, we demonstrated that human *TG* (length of 280 kb) exhibits TLs in thyrocytes of a freshly operated human thyroid (Fig. 2). Next, we detected the thyroglobulin gene (*tg*) in the chicken *Gallus domesticus* (length of ca.140 kb) and the frog *Xenopus tropicals* (length of ca.154 kb), verifying that the gene forms TLs in these species as well (Fig. 2). Finally, we aimed at visualization of the thyroglobulin gene in fish species, where thyrocytes are assembled into single follicles that are not gathered into a gland but remain separate and scattered along the esophagus. Therefore, to overcome difficulties of finding follicles, we made use of the transgenic zebrafish *Danio rerio* lines generated previously, in which thyrocytes express fluorophores under the *tg* promoter, tg(*tg*:mCherry) or tg(*tg*:GFP) (Opitz et al., 2013; Opitz et al., 2012). Using cryosections from these transgenic lines, we demonstrated that, despite the relatively small length of the *D*.*rerio tg* (68 kb), it forms microscopically resolvable TLs strongly expanding throughout the nucleus (Fig. 2).

**Figure 2.**
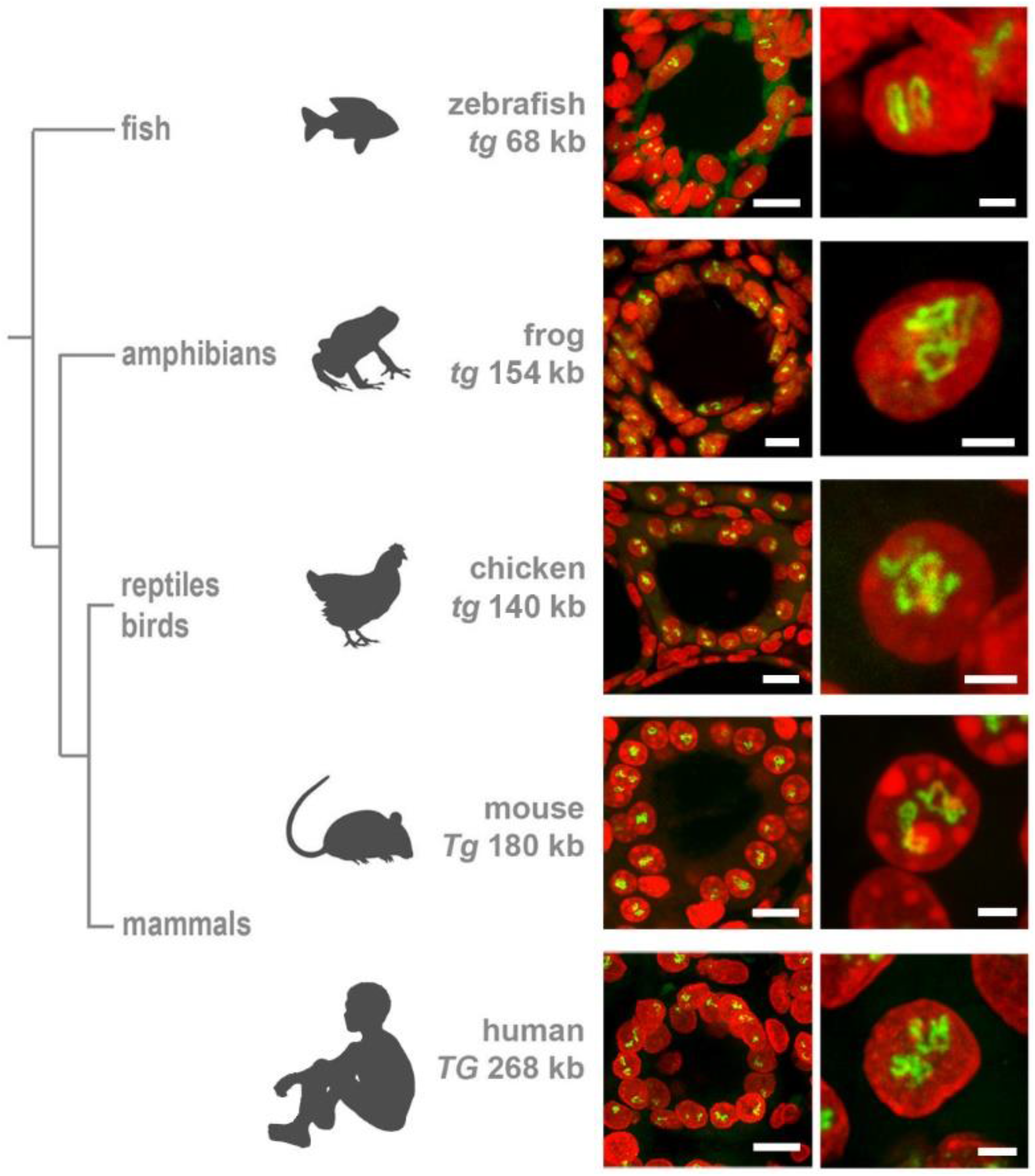
Thyroglobulin TLs are a peculiarity of thyrocytes from all major vertebrate groups. RNA-FISH with species-specific probes hybridizing to nascent RNA transcripts of thyroglobulin genes reveals TLs in all studied species. *The left column* shows thyroid follicles; *the right column* shows representative thyrocyte nuclei at a higher magnification. Note that even the shortest among vertebrates, zebrafish *tg* with a length of only 68 kb, forms TLs resolvable by light microscopy. RNA-FISH signals (*green*), DAPI (*red*); images are projections of confocal *3-6 µm* stacks. Scale bars: the left column, *10 µm*; the right column, *2 µm*.

Importantly, all the characteristic TL features, shown previously only for mouse TLs, are utterly manifested in other species as well. In particular, we observed separation of TL flanks caused by the increased gene stiffness (Fig. 3A,B), as well as co-transcriptional splicing detected by hybridization of sequential probes highlighting introns (Fig. 3C,D).

**Figure 3.**
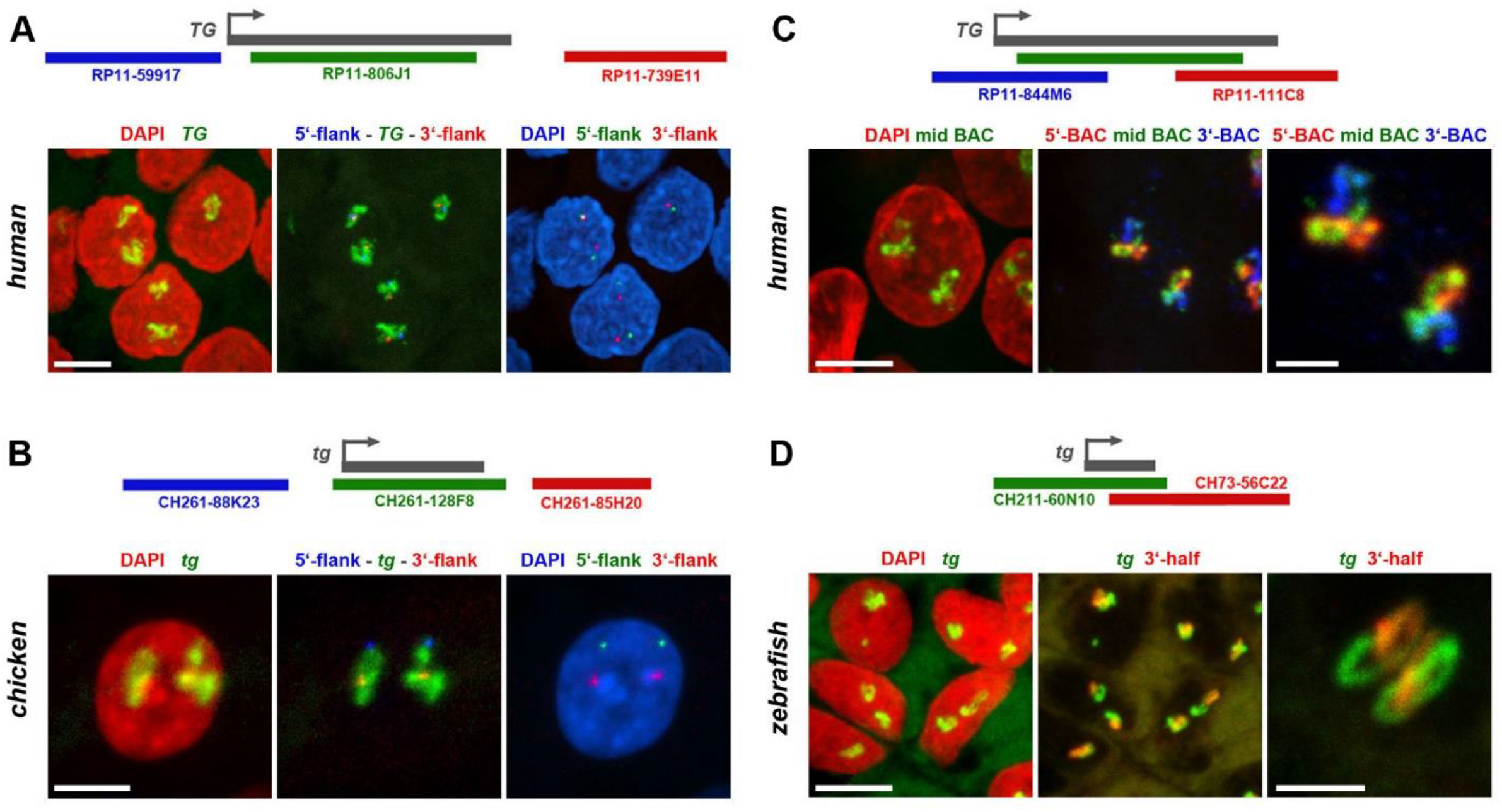
Thyroglobulin TLs in other vertebrates exhibit the key features of TLs. **(A**,**B)** As exemplified by simultaneous RNA- and DNA-FISH for human (A) and chicken (B) thyroglobulin, their TLs are open loops with visibly separated flanks. Images on the *left* show thyrocyte nuclei (*red*) with entire TLs (*green*). *Mid* images show the same TLs in *green* and the two flanks in *blue* (5’) and *red* (3’). Images on the *right*, for better demonstration of flank separation, show the same nuclei (blue) and the two flanks in *green* (5’) and *red* (3’). **(C**,**D)** As exemplified by RNA-FISH for human (C) and zebrafish (D) thyroglobulin genes, genomic probes highlighting introns, label TLs sequentially as a result of a co-transcriptional splicing. *Left* images show thyrocyte nuclei (*red*) with entire TLs (*green*). *Mid* images show the same TLs after hybridization with mid (*green)*, 5’ (*blue*) and 3’ (*red*) BACs (C). For zebrafish *tg*, only two BACs were used, one covering the whole gene (*green*) and one – its 3’ half (*red*) (D). Images on the *right* show TLs at a higher magnification with the same colour code. Schematics of genes and used BACs are shown above every panel. Images are projections of confocal stacks through *6 µm* (A,C), *3*.*5 µm* (B) and *2*.*5 µm* (D). Scale bars: A-D, *5 µm*; right panels on C and B, *2 µm*.

These data show a high upregulation of the thyroglobulin gene in the major vertebrate classes confirming the key role of TG in hormone production. This conclusion is reinforced by the recent work demonstrating a highly conserved structure of TG protein among vertebrates with all domains conserved from basal vertebrates to mammals (Di Jeso and Arvan, 2016; Holzer et al., 2016). Moreover, the authors of this paper conclude that since no orthologues of thyroglobulin gene have been found in invertebrates, TG protein and the way of THs generation are the vertebrate inventions.

### *Tg* TLs are a robust mark of thyrocytes differentiated *in vitro*

Sabine Costagliola’ s research group described the generation of a functional thyroid tissue *in vitro* in 2012 (Antonica et al., 2012). The authors demonstrated that transient overexpression of the transcription factors Nkx2-1 and Pax8 in mouse embryonic stem cells, followed by 3D culturing, allows the generation of follicles similar in morphology and gene expression to the thyroid follicles *in vivo*. 10 years later, the same group using similar strategy succeeded in generating and growing human thyroid organoids (Romitti et al., 2022). To confirm the functionality of mouse and human follicles generated *in vitro*, cultured follicles of both species were grafted into kidneys of athyroid mice resulting in successful rescue of hormone production after 4-5 weeks.

Not surprisingly, RNA-FISH revealed *Tg* TLs in mouse and human follicles cultured *in vitro*, although the loop size was noticeably smaller in comparison to thyrocytes in glands (Fig. S2). To visualize mouse *Tg* TLs in the grafted thyrocytes, we used RNA- and DNA-FISH simultaneously - to identify injected male mouse cells in the female host tissue by a probe for the Y chromosome (DNA-FISH) and to detect *Tg* loops (RNA-FISH). Accumulation of small follicles were clearly distinguished from the host kidney tissue by Y chromosome FISH signals (Fig. 4A), and predictably, only cells with this marker exhibited *Tg* TLs, which were much more extended compared to cultured follicles or even thyrocytes within the thyroid gland (Fig. 4A2,A3).

**Figure 4.**
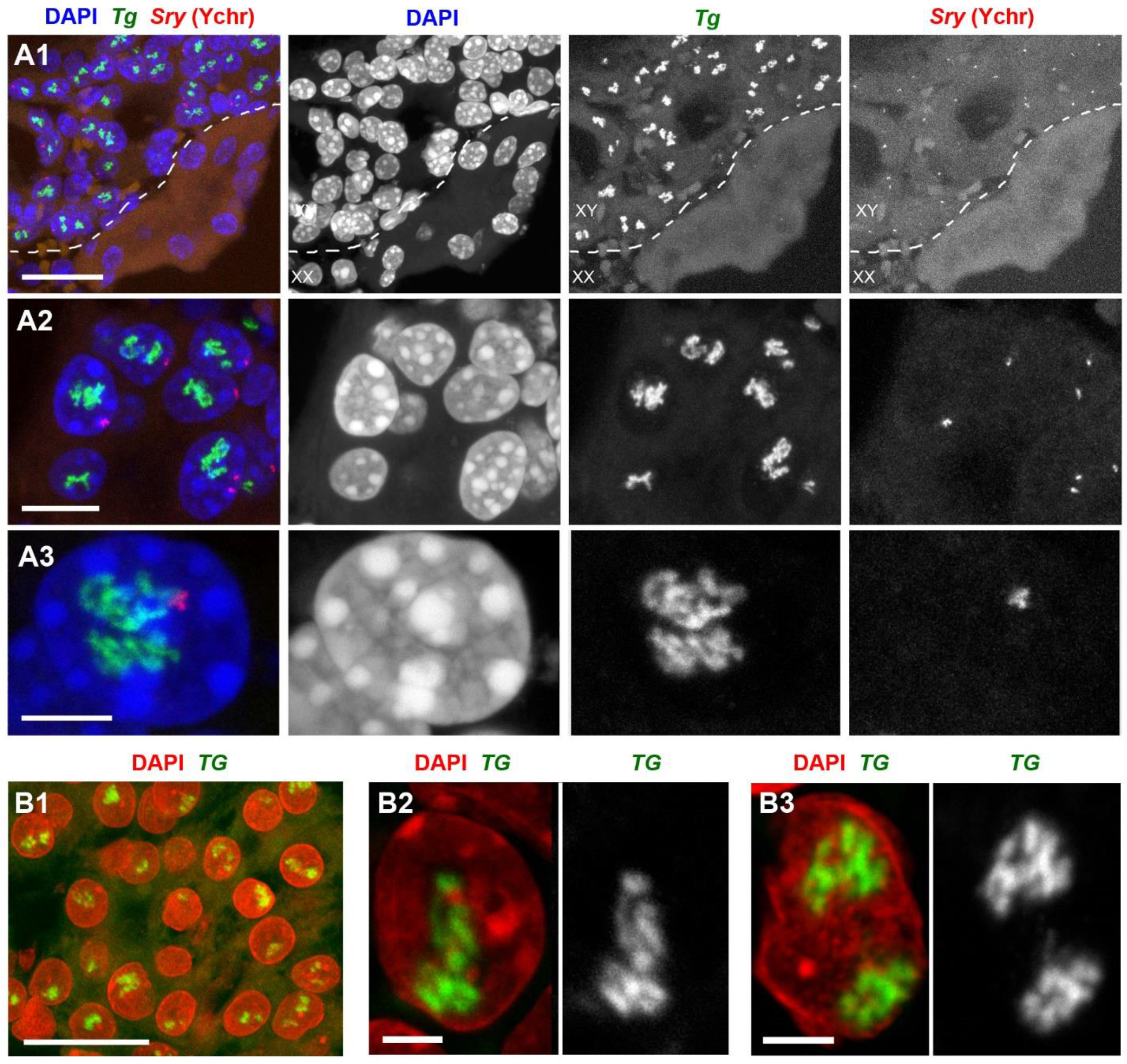
Mouse (A) and human (B) thyroid organoids grafted into mouse kidney. Cells of grafted follicles, generated from mouse male ESCs, can be distinguished from female mouse host cells by DNA-FISH with a Y chromosome-specific probe (*red*). The dotted line in **A1** separates host kidney cells from thyrocytes of follicles. Note the high extension of *Tg* and *TG* TLs in comparison to TLs formed by these genes in cultured follicles (compare to **Figure S2**) Thyroglobulin TLs are *green*; DAPI is *blue* in A and *red* in B panels. Images are projections of confocal *4 µm* stacks (A1 and B1) and *2*.*5 µm* (A2, A3, B2, B3). Scale bars: A1, B, *25 µm*, A2, *10 µm*, A3, *5 µm*, B2, B3, *2*.*5 µm*.

Differentiation between host mouse cells and human cells grafted into mouse kidney is an easy task, because in difference to human, mouse nuclei possess multiple chromocenters formed by subcentromeric major satellite repeat (Erdel et al., 2020; Vissel and Choo, 1989) brightly stained with DAPI (Fig. S3A). In addition, transgenic differentiated thyrocytes express NKX2-GFP and thus could be distinguished from non-thyrocytes (Fig. S3B). Similarly to mouse, human grafted thyrocytes form follicles (Fig. 4B1) and the *TG* genes become strongly expanded (Fig. 4B2,3). What is more, expressed *TG* modifies the harboring chromosomal locus by pushing its flanks away from each other (Fig. S3C) in a manner described for other resolvable TLs (Leidescher et al., 2022). Interestingly, using probes for the gene body and flanks, we noticed that several thyrocytes had four instead of two *TG* TLs (Fig. S3C), indicating that either during culturing *in vitro* or after grafting, some human thyrocytes became tetraploid. The *TG* TLs exhibit another typical TL feature, the co-transcriptional splicing: the three consecutive overlapping genomic probes sequentially label the nascent RNAs decorating the gene because the introns are sequentially spliced out (Fig. S3D). Thus, the work with thyrocytes differentiated from ESCs confirms that thyroglobulin TL formation is an invariable mark of differentiated thyrocytes.

### Thyrocytes are functional only in follicles

The noticeably stronger extension of both *Tg* and *TG* TLs in grafts in comparison to thyrocytes *in vitro* indicates a higher upregulation of the gene within an organism. Culturing of single thyrocytes isolated from thyroid confirms the importance of the follicle structure for thyrocyte functional activity. Following the protocol by (Jeker et al., 1999) for thyroid disintegration and culturing (see Methods), we generated primary transient cultures of mouse thyrocytes, consisting mostly of single thyrocytes and remnants of follicles. Immediately after disintegration, cells were attached to coverslips, fixed and hybridized with a *Tg* genomic probe. RNA-FISH showed that thyrocytes exhibit *Tg* TLs for at least an hour after follicle disintegration (Fig. S4A, B). After 24 h of incubation, however, single thyrocytes became flatter and lost *Tg* TLs (Fig. S4C), although strongly reduced TLs were still present in some cells within the remaining flattened follicles. After 72 h of incubation, all thyrocytes migrated out of follicles, became very flat and formed a monolayer, in which not a single cell exhibited *Tg* TLs (Fig. S4D). Apparently, thyrocytes lost their identity, possibly de-differentiated and entered the cell cycle with a high proliferative activity, as reported earlier for thyrocyte cultures established from other vertebrates (Kimura et al., 2001). These data indicate that results of various analyses conducted on cultured thyrocytes, as well as on other differentiated cells transferred to *in vitro* conditions, must be treated cautiously.

### *Tg* expression is independent of thyroidal TH status

TH production is tightly regulated by the activity of the hypothalamus-pituitary-thyroid axis and controlled by negative feedback loops involving the TH receptor THRB (Ortiga-Carvalho et al., 2016). Hypothalamic thyrotropin-releasing hormone (TRH) activates its pituitary TRHR1 receptor and stimulates the thyroid by stimulating hormone (TSH) release, which in turn acts on TSHR of thyrocytes and stimulates the production and secretion of THs. (Fig.5A). However, data on the regulation of the *Tg* gene activity are controversial. Earlier works showed that continuous presence of TSH is required to maintain TG production (Kim and Arvan, 1993; Van Heuverswyn et al., 1984). More recent work, however, showed that TSH deprivation or lack of functional TSHR due to early development does not affect the *Tg* expression but greatly reduces the expression of thyroperoxidase and the sodium/iodide symporter (Postiglione et al., 2002).

To test whether *Tg* expression is regulated by the TH status, we sampled thyroids from mice with increased or decreased thyroidal TH production further referred to as hyper- and hypothyroid conditions, respectively. For a hyperthyroid background we investigated *Mct8*-KO mice that exhibit a highly increased thyroidal TH content and reduced thyroidal T4 secretion (Trajkovic-Arsic et al., 2010; Trajkovic et al., 2007) (Fig. 5A,C) as well as *Thrb*-KO mice that show highly elevated TSH and TH levels (Forrest et al., 1996) (Fig. 5A,D). We also examined *Trhr1*-KO mice that display central hypothyroidism with decreased thyroidal and serum TH concentrations (Groba et al., 2013; Rabeler et al., 2004) (Fig. 5A,B). For each condition, three thyroids were fixed for RNA-FISH and other three used for qPCR.

**Figure 5.**
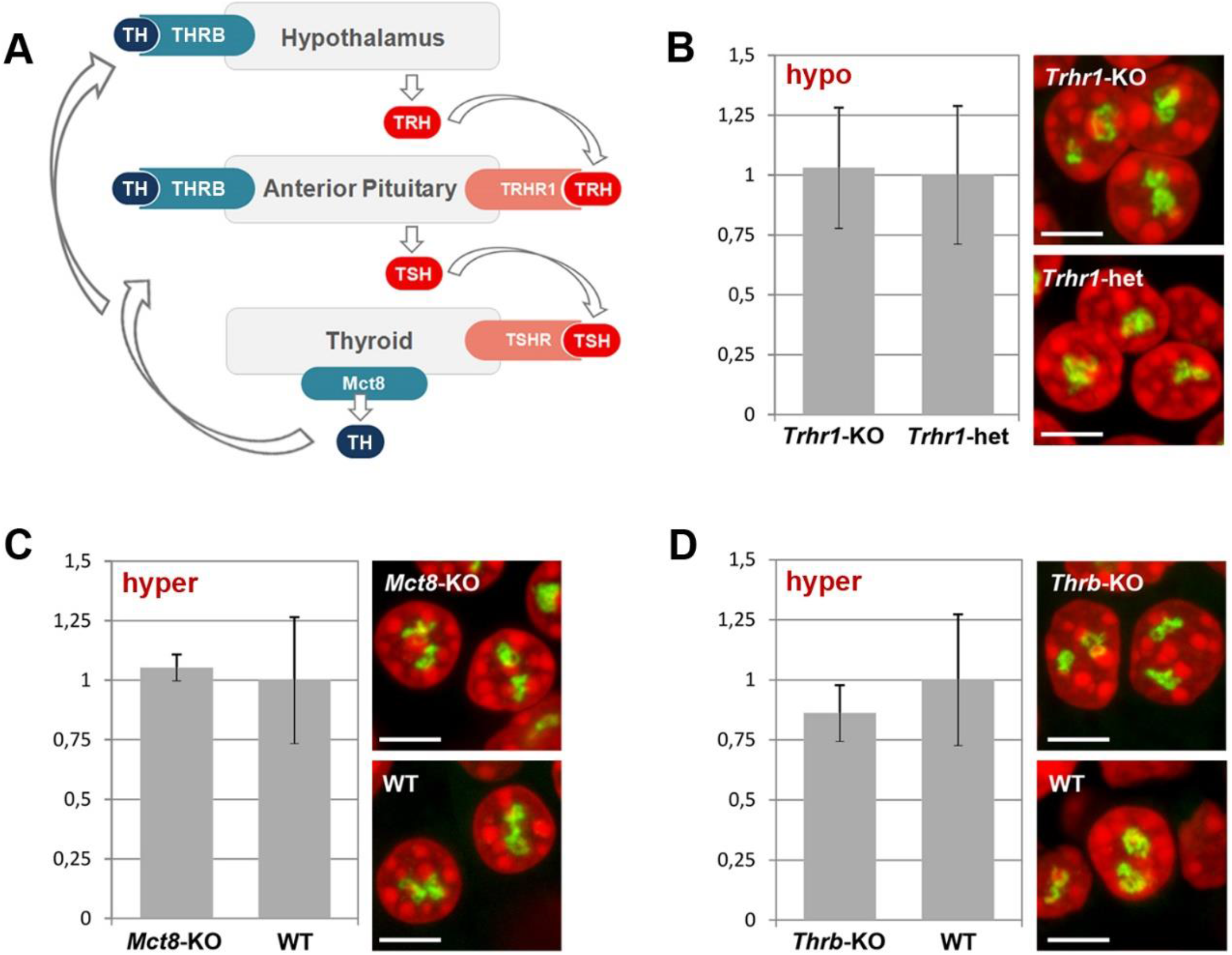
Thyroidal TH status does not influence *Tg* expression. **(A)** Simplified schematics of the hypothalamus-pituitary-thyroid axis. *TH*, thyroid hormones; *THRB*, TH receptor; *TRH*, hypothalamic thyrotropin-releasing hormone; *TRHR1*, thyrotropin-releasing hormone receptor; *TSH*, thyroid stimulating hormone; *TSHR*, thyroid stimulating hormone receptor. **(B-D)** Results of qPCR detecting levels of *Tg* expression (*left panels*) and RNA-FISH detecting *Tg* TLs (*right panels*) at a decreased (B) and increased (C-D) TH production in comparison to control mice. Bars in graphs are SDM. *Tg* TLs, *green*; DAPI, *red*; images are projections of *2-3 µm* confocal stacks; scale bars: *5 µm*.

We anticipated to observe a drop of *Tg* transcription level and consequently a decrease in *Tg* TL size under “hyper” conditions, whereas “hypo” condition with decreased thyroidal TH concentration might show elevated *Tg* transcription level and thus increased *Tg* TL size. In all three conditions, however, *Tg* mRNA level did not differ from control samples and thyrocytes exhibited *Tg* TLs of the size and morphology similar to those in control animals (Fig. 5B-D). Based on these results, we concluded that the intrathyroidal TH status does not influence *Tg* expression and that the gene is perpetually upregulated regardless of the activity of the hypothalamus-pituitary-thyroid axis.

### *Tg* is upregulated during both the exocrine and endocrine activities of thyrocytes

The thyrocytes function as both exocrine and endocrine glands. On the apical side, a thyrocyte secretes proteins (e.g. TG, TPO) into the follicle cavity filled with colloid, and on the basolateral site it releases TH into the circulation (Fig. 6A). Surprisingly, the question whether these two phases of activity happen simultaneously or during separate time-windows remains open. One possibility to separate the phases would be a regulated oscillation in synthesis and excretion of TG, which is the major component of the follicle colloid, due to a potential *Tg* gene circadian rhythmicity. Indeed, human TSH exhibits a clear circadian rhythm with a peak between 2 and 4 AM (Russell et al., 2008). This fact suggests that the *Tg* gene might also be subject to circadian activity, separating in this way the two thyrocyte physiological phases.

**Figure 6.**
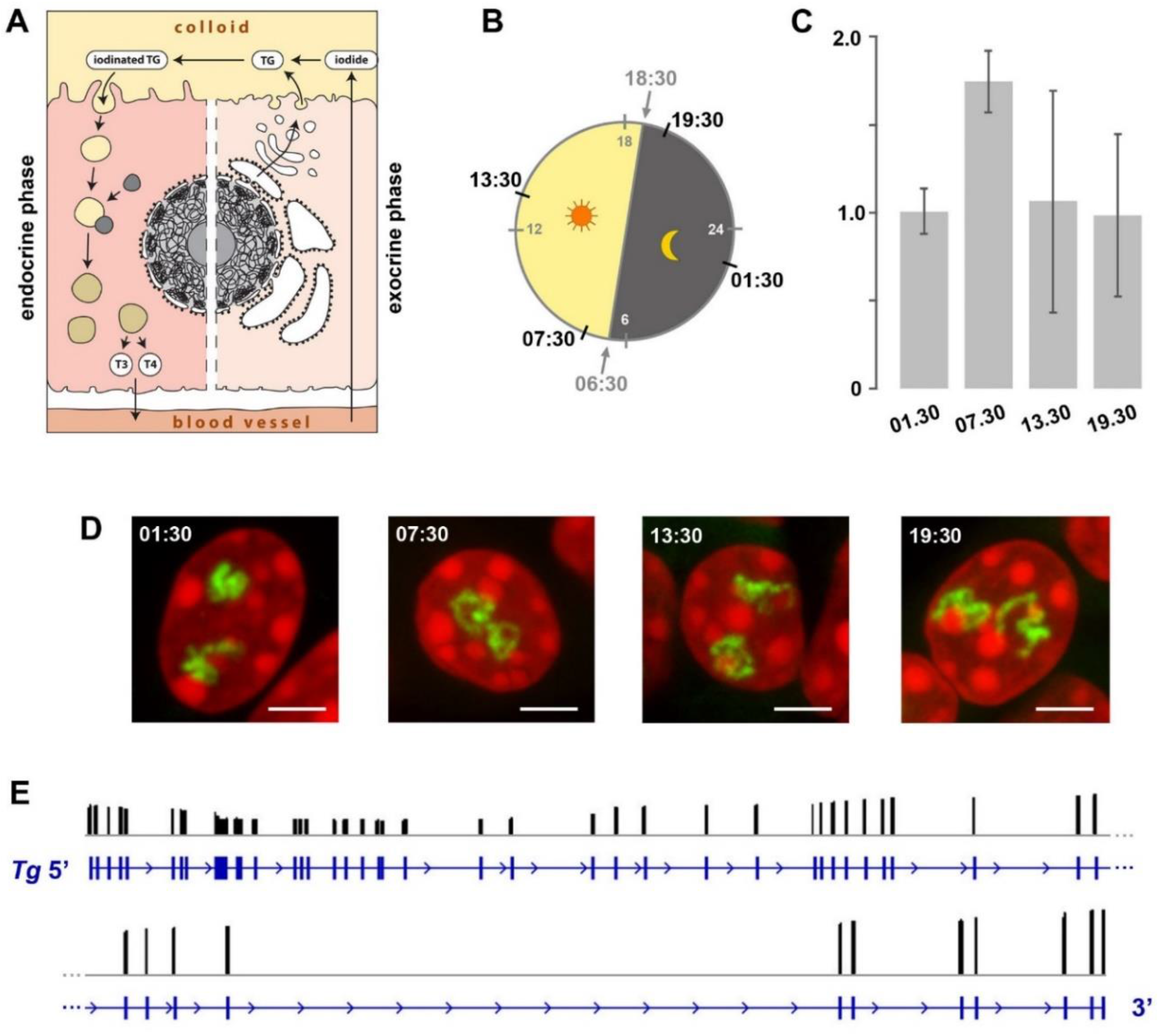
Expression of the *Tg* gene is not regulated by circadian activity and intron retention. **(A)** Schematics of a thyrocyte with two phases depicted - exocrine (*right*) and endocrine (*left*). **(B)** Diagram showing light : dark schedule (grey numbers) and time points (black numbers) of thyroid sampling for the circadian activity experiment. **(C)** Results of qPCR detecting levels of *Tg* expression indicate that there is no significant difference between the four time points. Bars are SDM. **(D)** Typical examples of thyrocyte nuclei after RNA-FISH detecting *Tg* TLs at four time points. *Tg* TLs, *green*; DAPI, *red*; images are projections of *3 µm* confocal stacks; scale bars: *5 µm*. **(E)** Results of Nano-pore sequencing of poly(A) fraction for the *Tg* gene. RNA-seq read coverage (*black columns*) across the gene locus (*blue*). The read coverage is ranging between 0 and 3146; only exon gene regions coincide with reads. For better presentation, the *Tg* gene is divided in two halves that are shown one under the other.

To check the possible oscillation of the *Tg* gene, we assessed the abundance of *Tg* RNAs and the presence of *Tg* transcription loops in thyroid glands from healthy mice round-the-clock. The mice were entrained to a 12 h : 12 h light : dark schedule and sacrificed at four time points according to one of the standard schemes for circadian activity testing. The time points included one hour after the light switched on (07:30), one hour after the light switched off (19:30), and two time points in the middle (13:30 and 01:30) (Fig. 6B). Thyroids of three animals were sampled for each time point, with one lobe of each thyroid used for qPCR to estimate *Tg* mRNA level and the other lobe used for microscopy. Visual inspection of *Tg* TLs after RNA-FISH did not reveal prominent differences in the sizes of the *Tg* TLs between three animals and four time points (Fig. 6D). In agreement with this, levels of *Tg* mRNA were also not significantly different (Fig. 6C). According to the above data, we conclude that the *Tg* gene does not exhibit rhythmic expression and remains highly upregulated throughout 24 hours.

The presence of extended *Tg* TLs in thyrocytes round-the-clock still leaves an opportunity for the regulation of gene expression by accumulation of transcripts in the nucleoplasm followed by their concurrent export into the cytoplasm. Such mechanism of fine gene expression tuning is known as the intron retention phenomenon described for various cell types (Braunschweig et al., 2014). Intron retention allows an acute release of mRNA into the cytoplasm by synchronous excision of retained introns in response to a stimulus (Mauger et al., 2016). Thus, despite the perpetual transcription of *Tg*, intron retention might be a mechanism for *Tg* mRNA accumulation within nuclei during the endocrine phase and their release during the exocrine phase. To test this hypothesis, we isolated RNA from three mouse thyroid glands and performed Nanopore sequencing. As shown in Fig. 6E, we have not detected intron retention in the poly(A) fraction of RNA: all sequenced reads mapped exclusively to the gene exons, which indicates that introns of all *Tg* mRNAs are excised. Therefore, we can rule out intron retention as a mechanism for restricting the TG production window.

## CONCLUSIONS

Collectively, our data suggest that both thyroglobulin alleles are perpetually upregulated in thyrocytes of the major vertebrate groups with a seemingly everlasting expression during the entire cell life. On one hand, it underscores the physiological importance of TH for proper organismal development and function. On the other hand, in view of the large volumes of TG stored within follicles, such high and enduring expression of the *Tg* gene, at any condition, at any age and round-the-clock seems counterintuitive and remains enigmatic.

Conceivably, it might reflect an inefficient way of hormone production evolved during vertebrate evolution. Indeed, only a small proportion of the 66 tyrosine residues of the thyroglobulin molecule becomes iodinated and only three or four TH molecules result from cleavage of one thyroglobulin molecule during thyrocyte endocrine (Di Jeso and Arvan, 2016; van de Graaf et al., 2001). Therefore, such wasteful thyroglobulin production might be the way to correct this unintelligible nature design. Another not mutually exclusive explanation of the phenomenon is that massive TG production is needed for storage of the rare trace element iodine (Crockford, 2009). One can speculate that binding to a large protein is a safe way of building an iodine reservoir within an organism.

Taking in account the very low turnover of thyrocytes (Dumont et al., 1992), we deduce that the thyroglobulin gene is perpetually active, e.g., for months in mouse and for years in human (Coclet et al., 1989). In this respect, the phenomenon of the thyroglobulin gene represents an attractive model to study transcription regulation, in particular, molecular mechanisms of high upregulation maintenance, chromatin dynamics, and kinetics of splicing.

## MATERIAL AND METHODS

### Tissue collection

All mouse studies were executed in accordance with the European Union (EU) directive 2010/63/EU on the protection of animals used for scientific purposes and in compliance with regulations by the respective local Animal Welfare Committees (LMU; Committee on Animal Health and Care of the local governmental body of the state of Upper Bavaria; Germany; Animal Welfare Committee of the Landesamt für Natur, Umwelt und Verbraucherschutz Nordrhein-Westfalen (LANUV; Recklinghausen, Germany). CD-1 mice were purchased from Charles River Laboratories, housed in individual cages with free access to food and water on a 12:12 light dark cycle at the Biocenter, Ludwig-Maximilians-University of Munich (LMU). Mice were sacrificed by cervical dislocation after IsoFlo (Isofluran, Abbott) narcosis. *Mct8*-KO mice (Trajkovic et al., 2007), *Trhr1*-KO mice (Rabeler et al., 2004); and *Thrb*-KO mice (Forrest et al., 1996), all on C57BL/6 background, were kept at 22°C in IVC cages in the central animal facility of the University Hospital Essen and had access to normal chow and water *ad libitum*. Adult mutant mice and control littermates were killed by CO2 inhalation prior to organ collection.

Human thyroid tissue was sampled from a freshly operated struma according to the ethical consent (ethical approval No. 12-5133-BO to DF) of the Department of Endocrinology, University Hospital Essen, Germany.

Thyroids from chicken (*Gallus gallus domesticus*) were collected from white legorn birds. Fertilized eggs of the M11 chicken line were kindly provided by Dr. S. Weigend (Federal Research Institute for Animal Health, Mariensee) and hatched at the Faculty for Veterinary Medicine, Munich. Birds were housed under conventional conditions in aviaries with groups of up to 10 birds and received food and water ad libitum. The animals were treated according to the standard protocol approved by The Committee on Animal Health and Care of the local governmental body of the state of Upper Bavaria, Germany. Thyroids from *Xenopus tropicalis* (*Silurana tropicalis*) were collected from four-year-old adult animals. Frogs were purchased at the Centre de Ressources Biologiques de Rennes (CRB) and raised in the PhyMa Laboratory in aquatic housing system (MPAquarien, Rockenhausen, Germany) at 24°C. Generation of transgenic zebrafish (*Danio rerio*) lines expressing fluorophores in thyrocytes, tg(tg:mCherry) and tg(tg:GFP) is described elsewhere (Opitz et al., 2013; Opitz et al., 2012).

In each case, freshly dissected tissues were washed with PBS and then fixed with 4% paraformaldehyde (Carl Roth) solution in PBS for 12-20 h.

### Mouse and human organoids

Generation of functional mouse and human thyroid tissues *in vitro* is described in (Antonica et al., 2012; Romitti et al., 2022). In our experiments we used thyrocyte follicles formed *in vitro* in 3D matrigel culture and follicles formed after grafting differentiated *in vitro* thyrocytes into mouse kidney. All grafting experiments were performed in accordance with local Animal Ethics (Commission d’ Ethique du Bien-Être Animal (CEBEA) Faculté de Médecine ULB, Project CMMI-2020-01).

### Primary thyrocyte culture

For cultivating mouse primary thyrocytes in vitro, the protocol from Jeker et al. 1999 was adapted. Briefly, thyroid glands of two mice, were minced into pieces using micro scissors under binocular and transferred into 2 ml tube containing 200 U/ml collagenase and 1 U/ml dispase in DMEM/F12/GlutaMax medium. Incubation was performed in a shaking thermo-block at 37°C for 2 h and followed by mechanical disruption using a glass Pasteur pipetting. Then cells were centrifuged at 2000 rpm (358 g) for 5 min and resuspended in DMEM supplemented with 10% FCS and 3% Pen/Strep. Single thyrocytes and follicles were seeded on coated coverslips (pre-incubated with 1 µg/ml polylysine) and cultured for 30 min, 24 h or 72 h before fixation with 4% formaldehyde.

### Cryosections

After fixation, thyroids were washed with PBS, cryoprotected in a series of sucrose, and embedded in Tissue-Tek O.C.T. compound freezing medium (Sakura). Blocks were stored at - 80°C before cutting into 16-20 µm sections using a cryostat (Leica CM3050S). Cryosections were collected on Superfrost Plus slides (Thermo Scientific) and stored at - 80°C before use.

### Electron Microscopy

Mouse thyroid glands were fixed with 2% glutaraldehyde in 300 mOsm cacodylate buffer (75 mM cacodylate, 75 mM NaCl, 2 mM MgCl_2_) for 30 min, postfixed with 1% OsO4 in the same buffer for 1 h at room temperature. After washings in distilled water, samples were incubated in 1% aqueous solution of uranyl acetate (Serva) for 1 h at 4 °C, dehydrated in ethanol series and acetone, and embedded in Epon Resin. Thin sections (50-70 nm) were prepared using Reichert Ultracut, stained with Reynolds lead citrate and examined with a transmission electron microscope (JEM 100 SX, JEOL) at 60 kV.

### Gene expression analysis

Dissected thyroids were immediately placed in RNAlater solution and total RNA was isolated using the NucleoSpin RNA Kit (Macherey-Nagel) according to the manufacturer’ s instructions. RNA integrity was checked by separating RNA fragments on a 1% agarose gel. Only samples with a 28S:18S rRNA ratio of ∼ 2:1 and without genomic contamination and RNA degradation were used for downstream applications. 1 µg of total RNA was reverse transcribed according to the manufacturer’ s instructions using the High capacity cDNA reverse transcription Kit with random primers (Applied Biosystems) or Maxima H Minus Reverse Transcriptase (Thermo Scientific) with gene specific primers. qPCR was performed in technical and biological triplicates in 10 µl reactions using LightCycler 480 SYBR Green Master Mix (Roche) or Luna Universal qPCR Master Mix (New England Biolabs) according to the manufacturer’ s instructions.

### Nanopore sequencing

Freshly dissected thyroids were immediately placed in ice cold TriZol and homogenized using an ultra-turrax dispersing tool. Poly(A)+ transcripts were isolated using magnetic oligoT-beads (Lexogen). The sequencing library was created using the PCR-cDNA Sequencing Kit (PCB111.24, Oxford Nanopore) and sequenced on a PromethION P24 on a R9.4.1 flowcell. The sequencing data were basecalled using Guppy v6.4.6 and mapped to the mouse genome (mm10) using minimap2.

### FISH probes

BAC clones encompassing the thyroglobulin genes and flanking regions of different vertebrate species, as well as *Tshr, Slc16a2* and *Tpo* mouse genes, were selected using the UCSC genome browser (see the list of BACs in the Table1) and purchased from BACPAC Resources (Oakland children’ s hospital) as agar stabs (https://bacpacresources.org/). BACs were purified via standard alkaline lysis or the NucleoBond Xtra Midi Kit (Macherey-Nagel), followed by amplification with the GenomiPhi Kit (GE Healthcare) according to the manufacturer’ s instructions. Amplified BAC DNA was labeled with fluorophores using homemade conjugated fluorophore-dUTPs by nick translation (Cremer et al., 2008). Labeled BAC DNA was ethanol precipitated with 10-fold excess of Cot-1 (1 mg/ml; Invitrogen, 18440-016) and 50-fold excess of salmon sperm DNA (5 µg/µl; Sigma), pellet was dried in a SpeedVak, and dissolved in hybridization mixture containing 50% formamid, 1xSSC and 10% of dextran sulphate. Oligoprobe for the *Tg* mRNA was generated using SABER-FISH protocol and described in detail previously (Leidescher et al., 2022).

**Table 1.** BACs used in the study

|  | Latin name | genomic region | BAC # |
| --- | --- | --- | --- |
| mouse | *Mus musculus* | *Tg*, gene body | RP24-229C15 |
|  |  | *Tshr*, gene body | RP23-152C21 |
|  |  | *Slc16a2*, gene body | RP23-114F20 |
|  |  | *Tpo*, gene body | RP24-172O17 |
| human | *Homo sapiens* | *TG*, gene body | RP11-806J1 |
|  |  | *TG*, 5' half | RP11-844M6 |
|  |  | *TG*, 3' half | RP11-111C8 |
|  |  | *TG*, 5' flank | RP11-599I7 |
|  |  | *TG*, 3' flank | RP11-739E11 |
| chicken | *Gallus domesticus* | *tg*, gene body | CH261-128F8 |
|  |  | *tg*, 5' flank | CH261-88K23 |
|  |  | *tg*, 3' flank | CH261-85H20 |
| frog | *Xenopus tropicalis* | *tg*, gene body | CH216-364N2 |
|  |  | *tg*, gene body | CH216-26O19 |
| zebrafish | *Danio rerio* | *tg*, gene body | CH211-184D12 |
|  |  | *tg*, 3' 3/4 of the gene | CH73-56C22 |

### FISH and immunostaining

FISH on cryosections was performed as previously described (Solovei, 2010). For DNA-FISH, sections were treated with 50 µg/ml RNaseA at 37°C for 1 h. For DNA-FISH or FISH detecting DNA and RNA simultaneously, denaturation of both probe and sample DNA was carried out on a hot block at 80°C for 3 min. For only RNA-FISH, RNasing and denaturation steps were omitted. Probes were loaded on sections under small glass chambers and sealed with rubber cement (for detail, see (Solovei, 2010)). Hybridizations were carried out in a water bath at 37°C for 2 days. After hybridization, rubber cement and chambers were removed, slides with sections were washed with 2xSSC at 37°C, 3 × 30 min, and then with 0.1xSSC at 60°C 1 × 7 min. Hybridized SABER probes were detected by incubating with 1 μM fluorescently labeled detection oligonucleotides in PBS for 1 h at 37°C followed by washing with PBS for 10 min.

The primary antibody for cytoplasmic TG detection (1:50; rabbit anti-TG Abcam, ab 156008) and the secondary donkey anti-rabbit conjugated with Alexa 555 (1:250; Invitrogen, Cat# A31570) were diluted in blocking solution (PBS + 2% BSA + 0.1% Saponin + 0.1% TritonX100) and applied under glass chambers covering sections. Incubation with primary and secondary antibodies were carried overnight at RT, in between and after incubations, sections were washed with PBS + 0.05% TritonX100 warmed up to 37°C; 3 × 30 min. In all experiments, nuclei were counterstained with 2 µg/ml DAPI in PBS for 30 min and Vectashield (Vector) was used as an antifade mounting medium.

### Microscopy

Confocal image stacks were acquired using a TCS SP5 confocal microscope (Leica) using a Plan Apo 63/1.4 NA oil immersion objective and the Leica Application Suite Advanced Fluorescence (LAS AF) Software (Leica). Z step size was adjusted to an axial chromatic shift and typically was either 200 nm or 300 nm. XY pixel size varied from 20 to 120 nm. Axial chromatic shift correction, as well as building single grey-scale stacks, RGB-stacks, montages and maximum intensity projections was performed using ImageJ plugin StackGroom (Walter et al., 2006). The plugin is available upon request.

## ACKNOWLEDGEMENTS

We are grateful to Maria Carmo-Fonseca (Instituto de Medicina Molecular João Lobo Antunes, Faculdade de Medicina da Universidade de Lisboa) for fruitful discussions. We thank Conny Niemann and Andreas Klingl (Biozentrum, LMU, Munich) for technical assistance with electron microscopy.

## AUTHOR CONTRIBUTIONS

*Conceptualization:* I.S., H.H., S.C.; *Methodology:* S.U., S.L., Y.F.,K.T, JB.F., B.M., I.S., H.H., S.K., H.B., B.K.; *Validation:* S.U., S.L., Y.F., K.T, B.M., I.S., H.H., S.K., H.B.; *Formal analysis:* S.U., S.L.; *Investigation:* S.U., S.L., Y.F., B.M., I.S., H.H., S.K., H.B.; *Resources:* JB.F., B.K., M.R., F.W., D.F.; *Data curation:* I.S., H.H., S.C., H.L.; *Writing - original draft:* S.U., I.S., H.H., S.C., Y.F., H.L.; *Writing - review & editing*: S.U., I.S., H.H.; *Visualization*: S.U., S.L., I.S.; *Supervision:* I.S.; *Project administration*: I.S.; *Funding acquisition:* I.S., H.L.

## DATA AVAILABILITY

All relevant data can be found within the article and its supplementary information. Data on Nanopore sequencing are deposited to GEO with GSE233457 accession number.

## SUPLEMENTARY FIGURES

**Figure S1.**
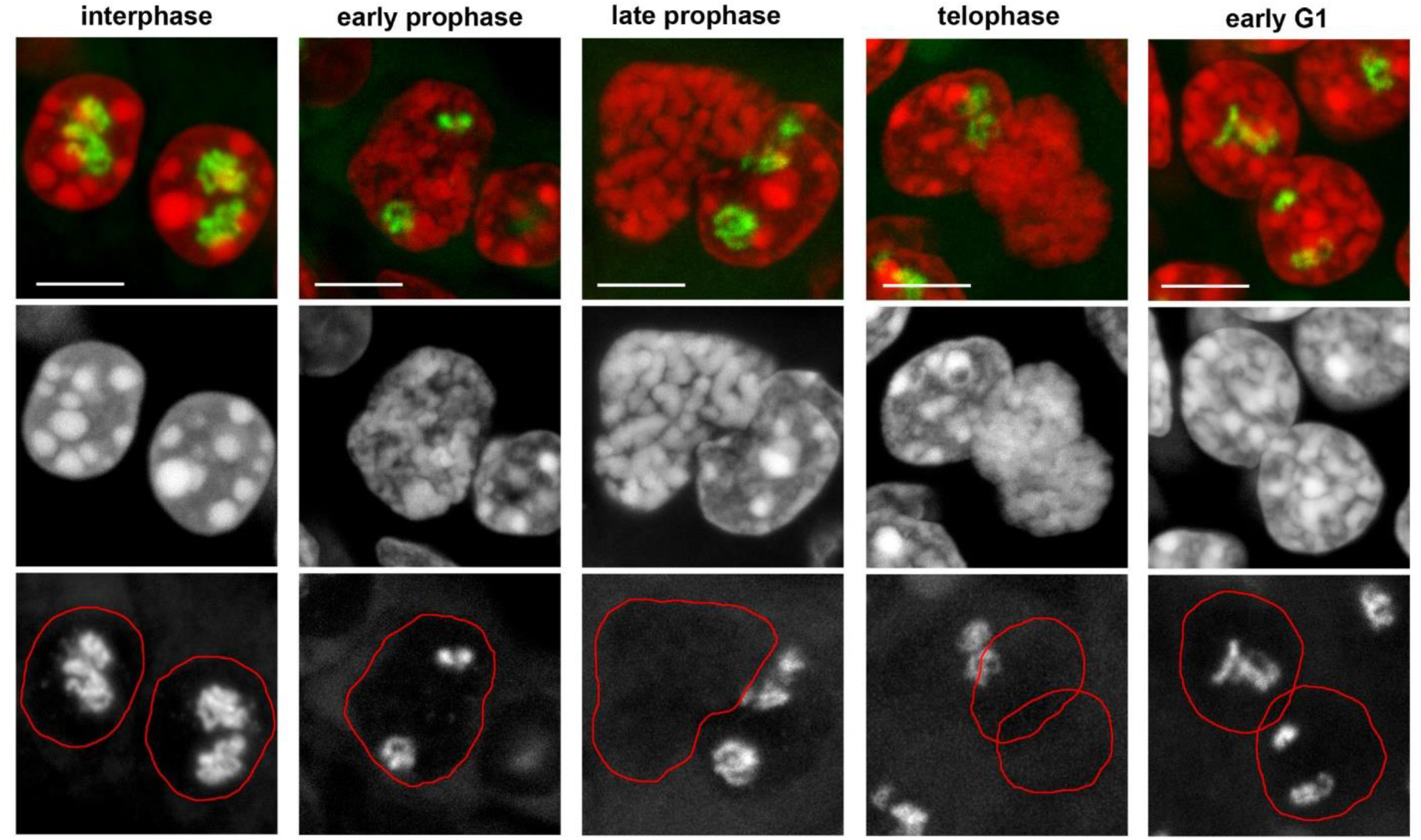
The *Tg* gene in cycling thyrocytes. TLs remain detectable by RNA-FISH until early-to-mid prophase, disappearing only during late prophase. The gene remains silent during metaphase, anaphase and early telophase, and becomes active in early G1. Note the *Tg* TLs are present when most chromatin is already (prophase) or still (G1) condensed. *Top panel* shows overlay of *Tg* signals (*green*) and DAPI (*red*); *mid panel* shows grey scale images of DAPI counterstain; *bottom panel* shows FISH signals with nuclear borders outlined with a *red line*. Images are projections of *3-6 µm* confocal stacks. Scale bars: *5 µm*.

**Figure S2.**
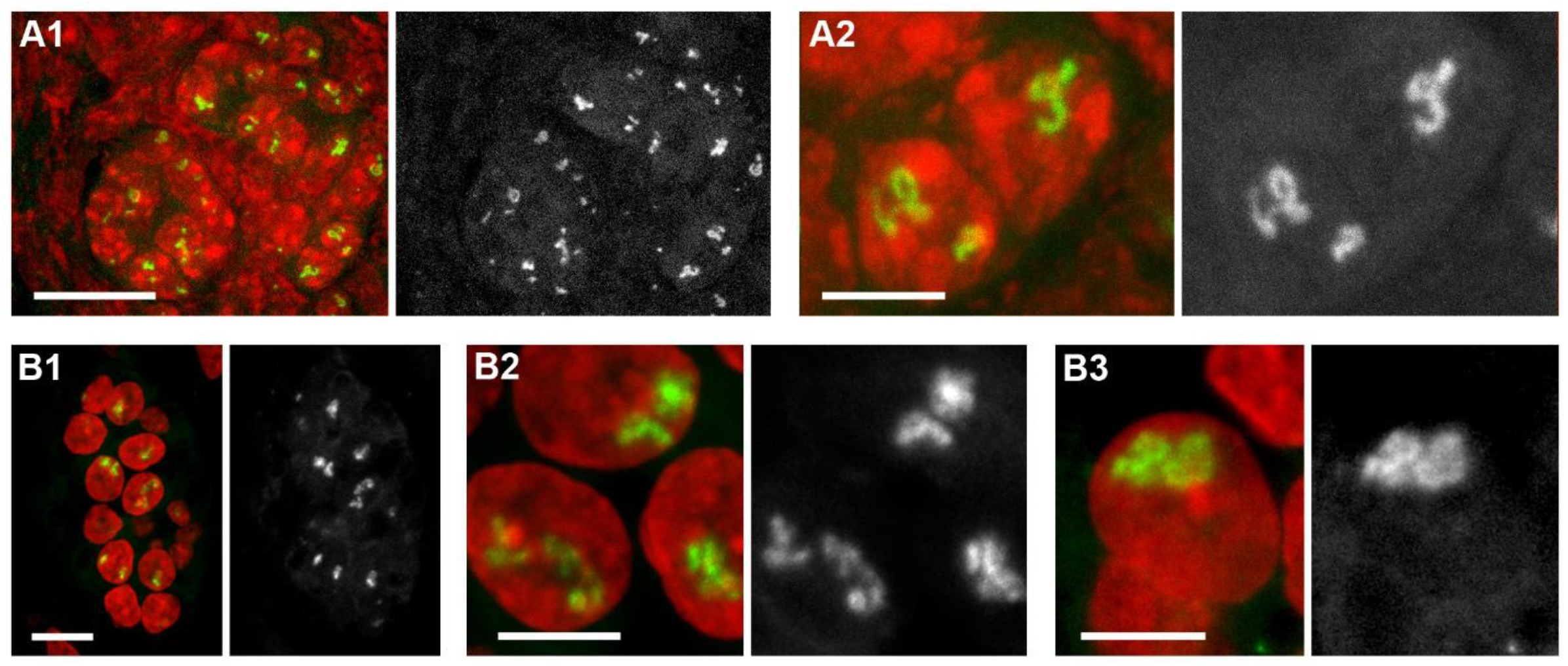
The *Tg* TLs in mouse (A) and human (B) organoids. RNA-FISH detecting *Tg* TLs (*green*) in nuclei (*red*) of thyroid organoids. Follicles formed in culture with matrigel (A1, B1) and thyrocytes within the follicles at a higher magnification (A2, B2, B3). For clarity, every RGB panel is supplemented by grey scale images of TLs. Images are projections of *4*.*5 µm* confocal stacks. Scale bars: A1,B1, *10 µm*, A2, B2, B3, *5* µm.

**Figure S3.**
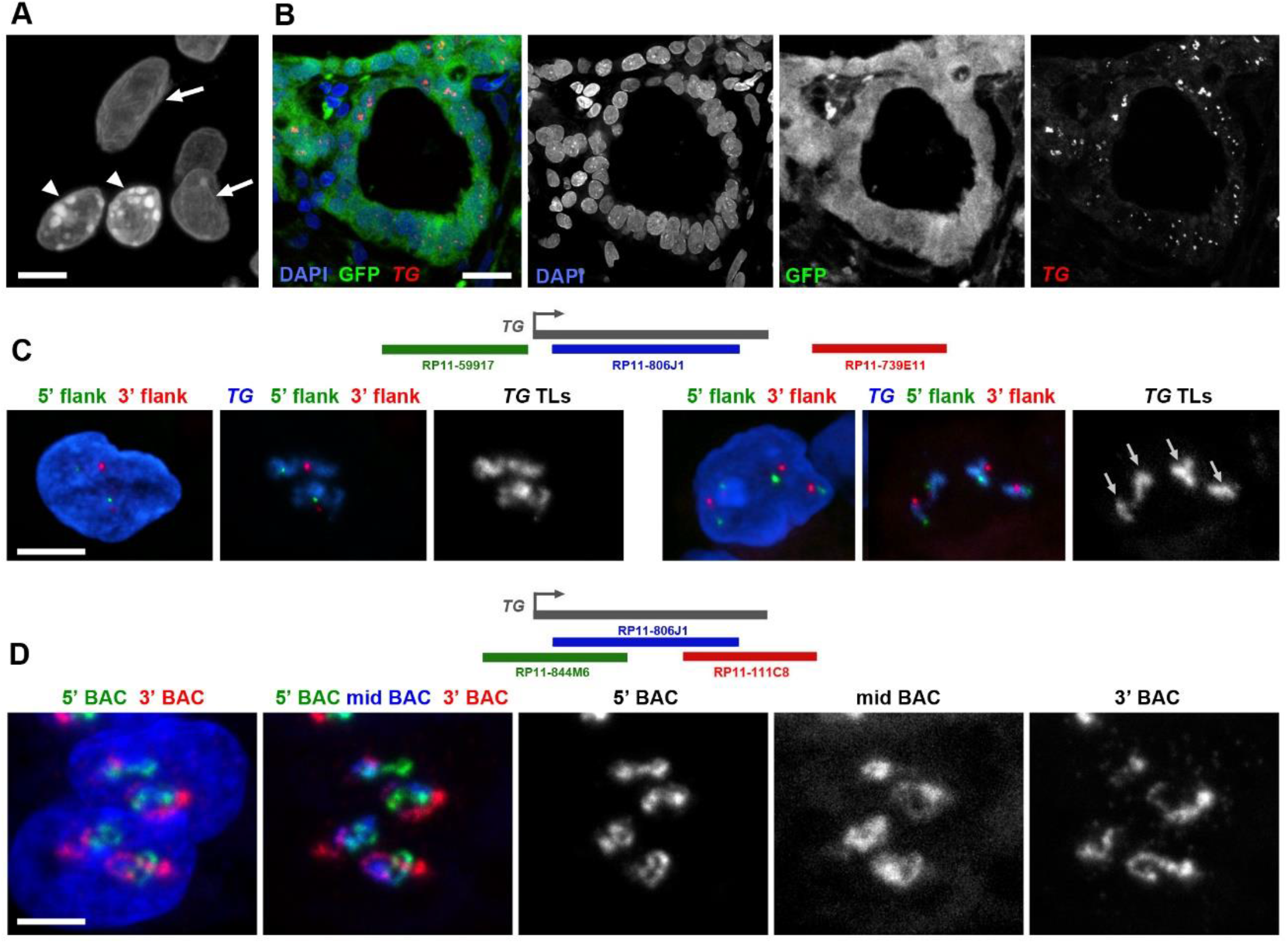
*TG* genes expressed in grafted follicles manifest typical TL features. **(A)** Human thyrocytes (*arrows*) grafted into mouse kidney can be unambiguously distinguished from mouse cells (*arrowheads*), because they possess brightly stained with DAPI chromocenters. **(B)** In addition, grafted human thyrocytes can be distinguished by the presence of GFP fluorescence (for detail, see Rommiti et al 2022). **(C)** Flanks of *TG* TLs are visibly separated. Left panels show grafted thyrocyte nuclei and 5’ (*green*) and 3’ (*red*) flanks; mid panels show flanks and TLs in *blue*; right panels show only TLs as grey scale images. Note that the nucleus on the right is from a tetraploid human thyrocyte with four *TG* TLs marked with arrows. **(D)** *TG* TLs are decorated by nRNA transcripts undergoing co-transcriptional splicing. Three BAC probes highlighting introns sequentially label *TG* loops. The left panel shows the nucleus in *blue*, the 5’ BAC in *green* and the 3’ BAC in *red*; the next RGB panel shows all three BACs including the middle one (*blue*). For clarity, the other three panels show grey scale images of each BAC signal. Schematics of genes and used BACs are shown above C and D panels. Nuclei are counterstained with DAPI (*blue*). Images are projections of *2-3 µm* confocal stacks. Scale bars: B, *25 µm*, A,C,D, *5 µm*.

**Figure S4.**
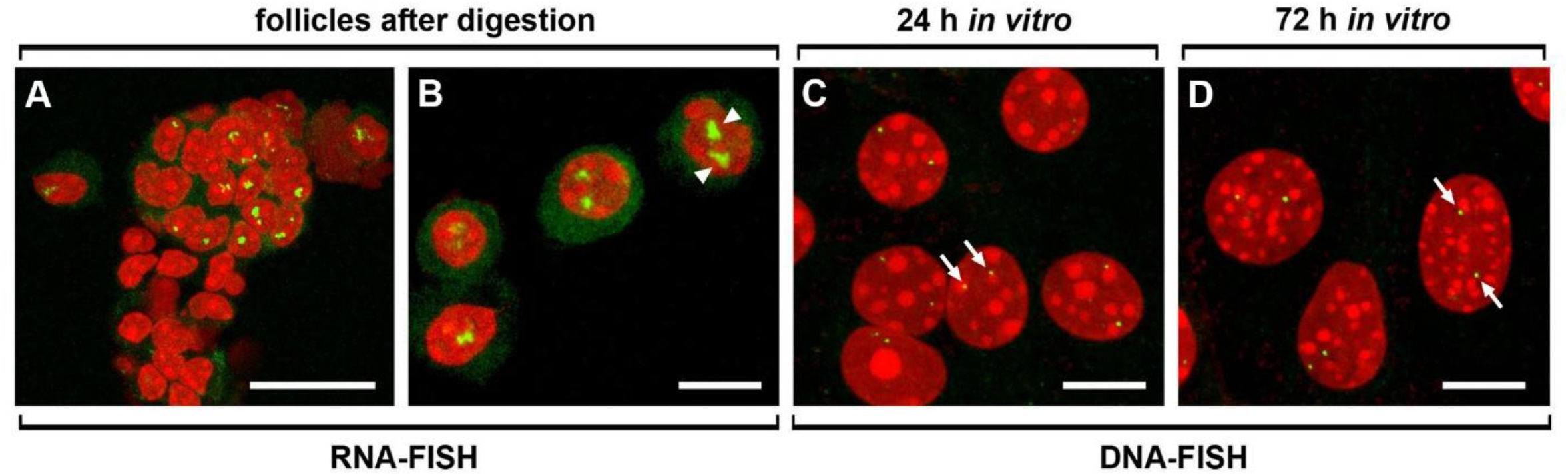
Withdrawal of *Tg* TLs in primary thyrocyte culture. **(A, B)** Freshly isolated follicle (A) and single thyrocytes (B) after RNA-FISH highlighting *Tg* TLs (*arrowheads*). **(C, D)** After culturing for 1 (C) and 3 (D) days, mostly cubical thyrocytes flatten, enter the cell cycle and withdraw *Tg* TLs. Since the *Tg* genes become silenced and condensed, they can be detected only by DNA-FISH producing dot-like signals (*arrows*). Note the difference in the nuclear morphology of freshly isolated (B) and cultured for 3 days (D) cells. DAPI, *red*; RNA and DNA signals, *green*; images are projections of *3-5 µm* confocal stacks; scale bars, *10 µm*

## REFERENCES

Antonica, F., D.F. Kasprzyk, R. Opitz, M. Iacovino, X.H. Liao, A.M. Dumitrescu, S. Refetoff, K. Peremans, M. Manto, M. Kyba, and S. Costagliola. 2012. Generation of functional thyroid from embryonic stem cells. Nature. 491:66–71.

Braunschweig, U., N.L. Barbosa-Morais, Q. Pan, E.N. Nachman, B. Alipanahi, T. Gonatopoulos-Pournatzis, B. Frey, M. Irimia, and B.J. Blencowe. 2014. Widespread intron retention in mammals functionally tunes transcriptomes. Genome Res. 24:1774–1786.

Coclet, J., F. Foureau, P. Ketelbant, P. Galand, and J.E. Dumont. 1989. Cell population kinetics in dog and human adult thyroid. Clin Endocrinol (Oxf). 31:655–665.

Cremer, M., F. Grasser, C. Lanctot, S. Muller, M. Neusser, R. Zinner, I. Solovei, and T. Cremer. 2008. Multicolor 3D fluorescence in situ hybridization for imaging interphase chromosomes. Methods Mol Biol. 463:205–239.

Crockford, S.J. 2009. Evolutionary roots of iodine and thyroid hormones in cell-cell signaling. Integr Comp Biol. 49:155–166.

De Felice, M., and R. Di Lauro. 2011. Minireview: Intrinsic and extrinsic factors in thyroid gland development: an update. Endocrinology. 152:2948–2956.

Di Jeso, B., and P. Arvan. 2016. Thyroglobulin From Molecular and Cellular Biology to Clinical Endocrinology. Endocr Rev. 37:2–36.

Dumont, J.E., F. Lamy, P. Roger, and C. Maenhaut. 1992. Physiological and pathological regulation of thyroid cell proliferation and differentiation by thyrotropin and other factors. Physiol Rev. 72:667–697.

Erdel, F., A. Rademacher, R. Vlijm, J. Tunnermann, L. Frank, R. Weinmann, E. Schweigert, K. Yserentant, J. Hummert, C. Bauer, S. Schumacher, A. Al Alwash, C. Normand, D.P. Herten, J. Engelhardt, and K. Rippe. 2020. Mouse Heterochromatin Adopts Digital Compaction States without Showing Hallmarks of HP1-Driven Liquid-Liquid Phase Separation. Mol Cell. 78:236–249 e237.

Forrest, D., E. Hanebuth, R.J. Smeyne, N. Everds, C.L. Stewart, J.M. Wehner, and T. Curran. 1996. Recessive resistance to thyroid hormone in mice lacking thyroid hormone receptor beta: evidence for tissue-specific modulation of receptor function. EMBO J. 15:3006–3015.

Groba, C., S. Mayerl, A.A. van Mullem, T.J. Visser, V.M. Darras, A.J. Habenicht, and H. Heuer. 2013. Hypothyroidism compromises hypothalamic leptin signaling in mice. Mol Endocrinol. 27:586–597.

Holzer, G., Y. Morishita, J.B. Fini, T. Lorin, B. Gillet, S. Hughes, M. Tohme, G. Deleage, B. Demeneix, P. Arvan, and V. Laudet. 2016. Thyroglobulin Represents a Novel Molecular Architecture of Vertebrates. J Biol Chem. 291:16553–16566.

Jeker, L.T., M. Hejazi, C.L. Burek, N.R. Rose, and P. Caturegli. 1999. Mouse thyroid primary culture. Biochem Biophys Res Commun. 257:511–515.

Kim, P.S., and P. Arvan. 1993. Hormonal regulation of thyroglobulin export from the endoplasmic reticulum of cultured thyrocytes. J Biol Chem. 268:4873–4879.

Kimura, T., A. Van Keymeulen, J. Golstein, A. Fusco, J.E. Dumont, and P.P. Roger. 2001. Regulation of thyroid cell proliferation by TSH and other factors: a critical evaluation of in vitro models. Endocr Rev. 22:631–656.

Leidescher, S., J. Ribisel, S. Ullrich, Y. Feodorova, E. Hildebrand, A. Galitsyna, S. Bultmann, S. Link, K. Thanisch, C. Mulholland, J. Dekker, H. Leonhardt, L. Mirny, and I. Solovei. 2022. Spatial organization of transcribed eukaryotic genes. Nat Cell Biol. 24:327–339.

Mauger, O., F. Lemoine, and P. Scheiffele. 2016. Targeted Intron Retention and Excision for Rapid Gene Regulation in Response to Neuronal Activity. Neuron. 92:1266–1278.

Mirny, L.A., and I. Solovei. 2021. Keeping chromatin in the loop(s). Nat Rev Mol Cell Biol. 22:439–440.

Opitz, R., F. Antonica, and S. Costagliola. 2013. New model systems to illuminate thyroid organogenesis. Part I: an update on the zebrafish toolbox. Eur Thyroid J. 2:229–242.

Opitz, R., E. Maquet, J. Huisken, F. Antonica, A. Trubiroha, G. Pottier, V. Janssens, and S. Costagliola. 2012. Transgenic zebrafish illuminate the dynamics of thyroid morphogenesis and its relationship to cardiovascular development. Dev Biol. 372:203–216.

Ortiga-Carvalho, T.M., M.I. Chiamolera, C.C. Pazos-Moura, and F.E. Wondisford. 2016. Hypothalamus-Pituitary-Thyroid Axis. Compr Physiol. 6:1387–1428.

Pauws, E., J.C. Moreno, M. Tijssen, F. Baas, J.J. de Vijlder, and C. Ris-Stalpers. 2000. Serial analysis of gene expression as a tool to assess the human thyroid expression profile and to identify novel thyroidal genes. J Clin Endocrinol Metab. 85:1923–1927.

Postiglione, M.P., R. Parlato, A. Rodriguez-Mallon, A. Rosica, P. Mithbaokar, M. Maresca, R.C. Marians, T.F. Davies, M.S. Zannini, M. De Felice, and R. Di Lauro. 2002. Role of the thyroid-stimulating hormone receptor signaling in development and differentiation of the thyroid gland. Proc Natl Acad Sci U S A. 99:15462–15467.

Rabeler, R., J. Mittag, L. Geffers, U. Ruther, M. Leitges, A.F. Parlow, T.J. Visser, and K. Bauer. 2004. Generation of thyrotropin-releasing hormone receptor 1-deficient mice as an animal model of central hypothyroidism. Mol Endocrinol. 18:1450–1460.

Romitti, M., A. Tourneur, B. de Faria da Fonseca, G. Doumont, P. Gillotay, X.H. Liao, S.E. Eski, G. Van Simaeys, L. Chomette, H. Lasolle, O. Monestier, D.F. Kasprzyk, V. Detours, S.P. Singh, S. Goldman, S. Refetoff, and S. Costagliola. 2022. Transplantable human thyroid organoids generated from embryonic stem cells to rescue hypothyroidism. Nat Commun. 13:7057.

Russell, W., R.F. Harrison, N. Smith, K. Darzy, S. Shalet, A.P. Weetman, and R.J. Ross. 2008. Free triiodothyronine has a distinct circadian rhythm that is delayed but parallels thyrotropin levels. J Clin Endocrinol Metab. 93:2300–2306.

Solovei, I. 2010. Fluorescence in situ hybridization (FISH) on tissue cryosections. In Methods Mol Biol. Vol. 659. 71–82.

Trajkovic-Arsic, M., J. Muller, V.M. Darras, C. Groba, S. Lee, D. Weih, K. Bauer, T.J. Visser, and H. Heuer. 2010. Impact of monocarboxylate transporter-8 deficiency on the hypothalamus-pituitary-thyroid axis in mice. Endocrinology. 151:5053–5062.

Trajkovic, M., T.J. Visser, J. Mittag, S. Horn, J. Lukas, V.M. Darras, G. Raivich, K. Bauer, and H. Heuer. 2007. Abnormal thyroid hormone metabolism in mice lacking the monocarboxylate transporter 8. J Clin Invest. 117:627–635.

van de Graaf, S.A., C. Ris-Stalpers, E. Pauws, F.M. Mendive, H.M. Targovnik, and J.J. de Vijlder. 2001. Up to date with human thyroglobulin. J Endocrinol. 170:307–321.

Van Heuverswyn, B., C. Streydio, H. Brocas, S. Refetoff, J. Dumont, and G. Vassart. 1984. Thyrotropin controls transcription of the thyroglobulin gene. Proc Natl Acad Sci U S A. 81:5941–5945.

Vissel, B., and K.H. Choo. 1989. Mouse major (gamma) satellite DNA is highly conserved and organized into extremely long tandem arrays: implications for recombination between nonhomologous chromosomes. Genomics. 5:407–414.

Walter, J., B. Joffe, A. Bolzer, H. Albiez, P.A. Benedetti, S. Muller, M.R. Speicher, T. Cremer, M. Cremer, and I. Solovei. 2006. Towards many colors in FISH on 3D-preserved interphase nuclei. Cytogenet Genome Res. 114:367–378.

